# Glenohumeral Injection Using Anatomic Landmark Versus Sonographic Needle Guidance

**DOI:** 10.1101/395293

**Authors:** Timothy S. Moore, Cole L. Paffett, Wilmer L. Sibbitt, William A. Hayward, James I. Gibb, Selma D. Kettwich, Roderick A. Fields, N. Suzanne Emil, Monthida Fangtham, Arthur D Bankhurst

**Affiliations:** Department of Internal Medicine, Division of Rheumatology, University of Pennsylvania, 3737 Market St Fl 8 Philadelphia, PA, USA 19104; Department of Surgery, Geisinger Health System, 100 N. Academy Ave, Danville, PA 17822; Department of Internal Medicine, Division of Rheumatology and School of Medicine, MSC 10 5550, 5th FL ACC, University of New Mexico Health Sciences Center, Albuquerque, NM, USA 87131.; The Department of Exercise and Sport Sciences, New Mexico Highlands University, Las Vegas, NM, USA; Department of Internal Medicine, Division of Nephrology, MSC 10 5550, 5th FL ACC, University of New Mexico Health Sciences Center, Albuquerque, NM, USA 87131.; 338 Amherst Dr. NE, Albuquerquerque, NM, 87106; Department of Internal Medicine, Division of Rheumatology and School of Medicine, MSC 10 5550, 5th FL ACC, University of New Mexico Health Sciences Center, Albuquerque, NM, USA 87131. Email

**Keywords:** Ultrasound, shoulder, glenohumeral, injection, corticosteroid, osteoarthritis

## Abstract

**Objective:** We hypothesized ultrasound (US) guidance improves outcomes of corticosteroid injection of the painful shoulder.

**Methods:** 30 patients with symptomatic shoulders due to osteoarthritis were randomized to glenohumeral injection with 3 milliliters of 1% lidocaine and 60 mg of triamcinolone acetonide using the anterior approach with 1) conventional anatomic landmark palpation-guidance or 2) US-guidance. Injection pain (visual analogue pain scale (VAS)), pain at outcome (2 weeks and 6 months), therapeutic duration, time-to-next-injection, and costs were determined.

**Results:** Injection pain was less with US (VAS: 0.3±0.6 cm) vs. landmark-guidance (VAS: 1.4±2.4 cm, 95% CI of difference: 0.5<1.1<1.7, p=0.05). Pain scores were similar at 2 weeks: US: 2.2±2.4 cm; Landmark: 1.8±2.7 cm, 95% CI of difference: −2.2<−0.4<1.4, p=0.66 and 6 months: US: 5.8±2.8 cm; Landmark: 6.4±2.9 cm, 95% CI of difference: −0.4<0.6< 1.1, p =0.71. Therapeutic duration (US: 3.9±1.5 months; Landmark: 3.0±1.2 months, 95% CI of difference: − 1.4 <−0.9<−0.4, p=0.045) and time-to-next-injection (US: 8.1±3.5 months; Landmark: 5.7±2.9 months, 95% CI of difference: −3.6<−2.4<−1.3, p=0.025) were longer, and fewer injections per year (29% less) were required: US: 1.5±0.2 injections/year; Landmark: 2.1±0.2 injections/year (p<0.037; 95% CI of difference −0.9<−0.6<−0.3). However, cost/patient/year was modestly greater with US (US: $318±89, Landmark: $301±67; p=0.28).

**Conclusion:** Anatomic landmark guidance in the short-term is equally effective as US for injection of the osteoarthritic shoulder and modestly less costly, however, US may reduce the need for repetitive injections by prolonging the therapeutic effect and thus time to next injection.

**IRB Statement:** This project was in compliance with the Helsinki Declaration, was approved by the Institutional Review Board (IRB) as ultrasound subset of a syringe safety trial (Human Research Review Committee approval 04-347), and was registered at C*linicalTrials.gov* (Clinical Trial Identifier NCT00651625). The subjects gave informed consent to participate prior to all studies and interventions. Patient confidentiality was protected according to the U.S. Health Insurance Portability and Accountability Act (HIPAA) and all data was de-identified.

## INTRODUCTION

Therapy for the painful shoulder with adhesive capsulitis, mild osteoarthritis, or impingement syndrome includes physical therapy, local measures, range of motion exercises, and intraarticular corticosteroid injections (1-4). In an effort to improve the accuracy and effectiveness of shoulder injections, sonographic needle guidance has been used with increasing frequency; however, data regarding cost-effectiveness are wanting (5-8). As Narzarian et al and Harkey et al and others note this paucity of cost-effectiveness research has resulted in challenges to musculoskeletal imaging by third party payers and has provoked skepticism and resistance to fully integrating sonography at the clinic level (9-12).

The present randomized controlled study preliminarily addressed whether sonographic needle guidance improved the clinical outcomes and cost-effectiveness of corticosteroid injection in the painful shoulder.

## Methods

### Subjects

This project was in compliance with the Helsinki Declaration, was approved by the Institutional Review Board (IRB) as ultrasound subset of a syringe safety trial (Human Research Review Committee approval 04-347), and was registered at C*linicalTrials.gov* (Clinical Trial Identifier NCT00651625). The subjects gave informed consent to participate prior to all studies and interventions. Patient confidentiality was protected according to the U.S. Health Insurance Portability and Accountability Act (HIPAA) and all data was de-identified. Inclusion criteria for participants with painful shoulder associated with osteoarthritis included: 1) pain with passive abduction/elevation of the shoulder; 2) pain to deep palpation lateral to the coracoid process; 3) nocturnal shoulder pain; 4) significant pain in the affected shoulder by 0-10 cm Visual Analogue Pain Sale (VAS) where VAS ≥ 5 cm; 5) failure of rest, range of motion exercises, and nonsteroidal anti-inflammatory drugs, 6) radiographs that demonstrated only mild to moderate (Grade I-II) osteoarthritis of the glenohumeral joint, and 7) the desire of the participant to have a corticosteroid injection. Patients who fulfilled all inclusion criteria were eligible to participate in this study. Exclusion criteria included 1) shoulder deformity, 2) complete rotator cuff tear, 3) the diagnosis of cervical spine nerve root impingement, 4) recent trauma, 5) hemorrhagic diathesis, 6) use of warfarin or other anticoagulants, 7) the presence of infection, or 8) previous corticosteroid injection into the shoulder in the preceding 6 months.

30 patients were randomized to either 1) glenohumeral injection guided by anatomic landmark palpation (landmark-guidance) (15 shoulders) or 2) sonographic guidance (15 shoulders). All injections and ultrasound examinations in both groups were performed by experienced traditional (anatomic landmark) and ultrasound-trained proceduralists. As this was a practical study meant to resemble clinical practice, neither the patient nor proceduralist was blinded to ultrasound versus palpation; thus, a “mock ultrasound” procedure was not performed in controls. “Mock ultrasound” deviates from normal practice and may induce a consistent procedural error because the shoulder is injected with one hand while the other hand operates the ultrasound transducer, when in normal practice the proceduralist uses 2 hands to perform a landmark procedure. The physician who determined post-procedural outcomes was blinded to the intervention (ultrasound vs. landmark). Study groups were similar in terms of gender, age, and baseline pain (Table 1). 100% of participants completed the study.

**Table 1.** Subject Characteristics and Procedural Pain

|  | **Landmark Guidance** | **Sonographic Guidance** |  |  |  |
| --- | --- | --- | --- | --- | --- |
| **Number of shoulders** | 15 | 15 | **Percent Difference** | **95% CI of difference** | **P Value** |
| **Age\*** | 53.6±15.8 | 57.3±12.9 | 6.8% | -14.0 <-3.7< 6.6 | 0.49 |
| **Female Gender\*\*** | 93.3% (14/15) | 93.3% (14/15) | 0% | Not applicable | 0.52 |
| **Pre-Procedure Baseline Pain (VAS Pain Scale)\*** | 7.9±1.3 | 8.0±1.8 | 1.3% | -1.2 <-0.1< 1.0 | 0.86 |
| **Pre-Procedure Baseline Pain (VAS Pain Scale)\*** | 7.9±1.3 | 8.0±1.8 | 1.3% | -1.2 <-0.1< 1.0 | 0.86 |
| **Needle Introduction Pain (VAS Pain Score)\*** | 3.4±3.4 cm | 2.5±1.8 cm | -26% | -1.2 <0.9< 3.0 | 0.18 |
| **Significant Needle Introduction Pain (VAS > 5 cm)\*\*** | 20% (3/15) | 0% (0/15) | -100% | Not applicable | 0.11 |
| **Injection Pain (10 cm VAS Pain Score)\*** | 1.4±2.4 cm | 0.3±0.6 cm | -79% | 0.5 <1.1< 1.7 | 0.05 |
| **Significant Injection Pain (VAS ≥ 5 cm)\*\*** | 6.7% (1/15) | 0% (0/15) | -100% | Not applicable | 0.48 |

\*Mean± Standard Deviation, one-tail t-test
\*\*Fisher Exact Test

### Outcome Measures

Participant pain was measured with the standardized and validated 0-10 cm Visual Analogue Pain Scale (VAS), where 0 cm = no pain and 10 cm = unbearable pain and 0.1 cm increments were permitted (13,14). Pain by VAS was determined 1) prior to the procedure (baseline pain), 2) during the insertion of the needle (procedural pain), 3) 2 weeks post procedure (pain at primary outcome), and 4) 6 months post procedure or directly before the next injection if less than 6 months (secondary outcome). A printed VAS was given to the patient to rate the pain before and during the procedures. The operating physicians recorded the pre-procedural and procedural pain. A VAS was given to the patient to take home after the procedure. The patient was then contacted by a physician blinded to treatment (ultrasound versus landmark) to obtain the VAS scores at 2 weeks and 6 months and the patient related the VAS scale to the inquirer while the patient looked at the VAS scale at home. Therapeutic duration (duration of therapeutic response or time to flare) was defined as the time interval in months when the shoulder became symptomatic (VAS ≥ 5 cm) after the injection. The patient emailed or called when the shoulder became symptomatic again. No patients were permitted a second injection of the shoulder prior to 3 months after the first injection; however, after 3 months the patient determined the timing of the next shoulder injection. If the shoulder remained asymptomatic at 6 months or longer, the duration was defined as 6 months. Time-to-next-intervention in this study corresponds to the time to the next injection and thus the time-to-next-intervention permits least cost determinations related to injection therapy (15-19). In the present series, the next intervention in each case was reinjection with corticosteroid. During the time of the study no patients underwent shoulder surgery or advanced imaging, but most received a second injection within the 12-month period. Time-to-next-intervention was determined up to 12 months after the initial injection and expressed in months. If the next injection occurred at a time greater than 12 months, the time-to next-intervention was defined as 12 months. The total number of corticosteroid injections per year was recorded for each individual.

Costs of the injection procedure in US dollars ($) were defined as those costs reimbursed by 2017 Medicare (United States) national rates for HCPC/CPT 20610 code for a large joint arthrocentesis for a physician office ($53.32/procedure), HCPC/CPT 9921315 code for minute outpatient encounter ($76.68), 60 mg triamcinolone acetonide ($13.00/procedure), and HCPC/CPT code 99213 for ultrasound guidance for needle procedures ($71.74) (20). With these costs in mind, an anatomic landmark injection costs $143/procedure, and an ultrasound guided injection cost $215/procedure. Yearly costs were calculated by multiplying the costs/procedure x 12 months divided by the months to time-to-next-intervention (time-to-the-next injection) (18- 20).

### Sonographic Needle Guidance

The injection procedure of the shoulder was performed in a standardized manner using the anterior approach in both the ultrasound and anatomic landmark groups (21-24). The one-needle two-syringe technique was used where 1) one needle is used for anesthesia and corticosteroid injection; 2) the first syringe is used to anesthetize the shoulder and perform arthrocentesis, and 3) the second syringe is used to inject the corticosteroid therapy (18,19).

In the palpation guided anatomic landmark technique with the forearm held relaxed in the patient’s lap, first the acromioclavicular joint and the coracoid process were palpated and marked with ink (Figure 1). The medial border of the humeral head was then palpated and marked. Chlorhexidine 2% was used for antisepsis. A 22 gauge 2 inch needle on a 3 ml syringe and filled with 3 ml of 1% lidocaine was introduced horizontally on the lateral side of the coracoid through the ligaments and joint capsule into the synovial joint with care not to forcefully inject into the glenoid labrum. Fox et al have demonstrated that inclusion of local anesthetic into shoulder injections improves post-procedure pain measures (25). Arthrocentesis was performed and joint effusion was aspirated, if present. Pain scores were recorded. The syringe was then detached from the needle while still in the glenohumeral joint and a 3 ml syringe prefilled with 60 mg triamcinolone acetonide suspension (Kenalog® 40) was attached to the indwelling needle, and the treatment was slowly injected. The needle was then extracted, firm pressure applied to the puncture site, and a sterile bandage was applied.

**Figure 1.**
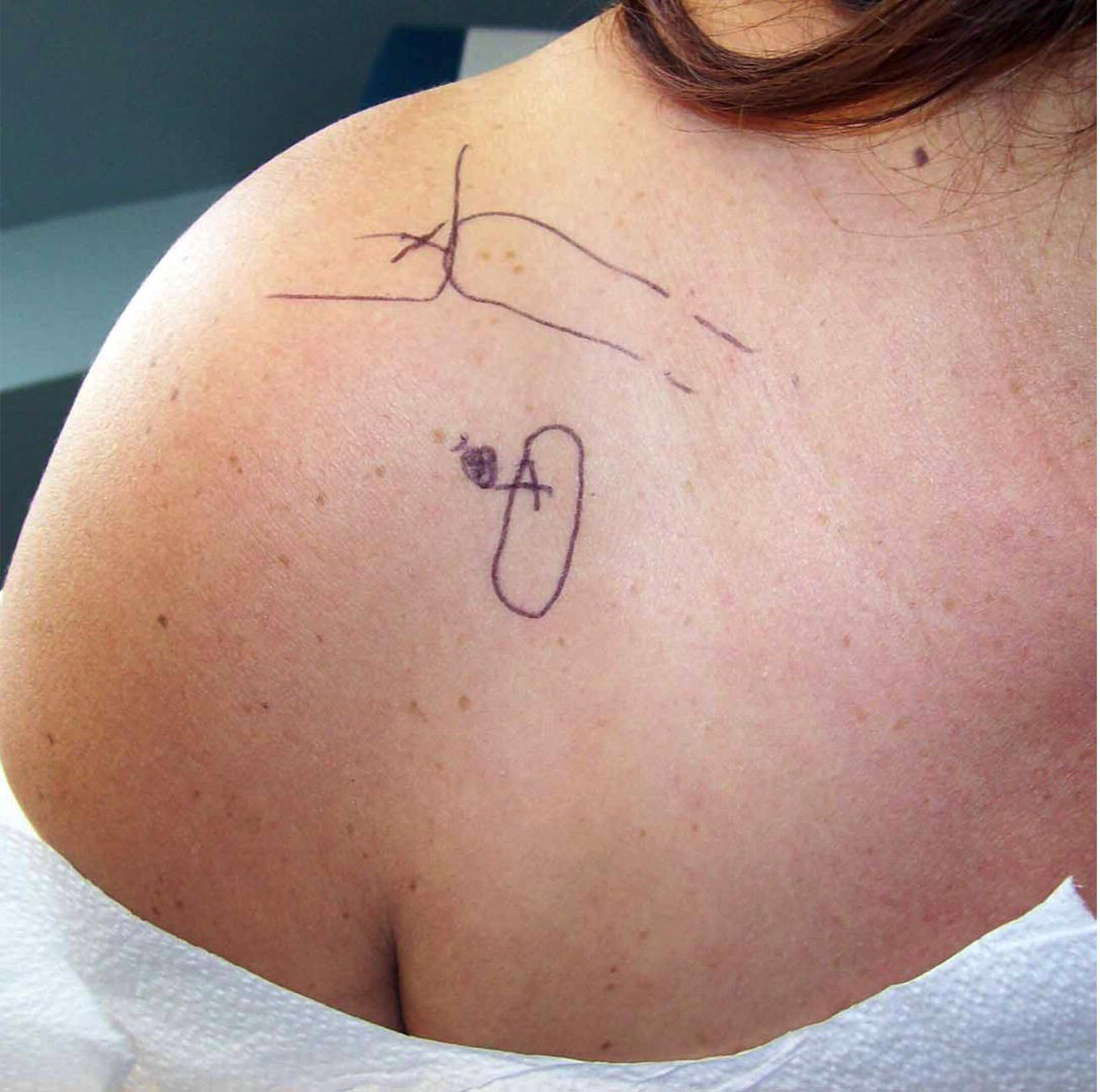
Anatomic landmark palpation anatomy for palpation-guided anterior glenohumeral joint injection. In the palpation guided anatomic landmark technique with the forearm held relaxed in the patient’s lap, first the acromioclavicular joint and the coracoid process are palpated and marked. The medial border of the humeral is then was then palpated and marked. The needle was introduced horizontally parallel to the floor on the lateral side of the coracoid through the ligaments and joint capsule into the glenohumeral joint with care not to forcefully inject into the glenoid labrum.

For the ultrasound-guided procedures a portable ultrasound unit with a 10-5 MHz 38 mm broadband liner array transducer (Sonosite M-Turbo, SonoSite, Inc. 21919 30th Drive SE, Bothell, WA 98021, website: www.sonosite.com) was used. In the ultrasound-guided technique the linear probe was held horizontally and the coracoid process, humeral head, and glenoid labrum were identified (Figure 2). The needle was then introduced out-of-plane inferiorly and laterally to the coracoid and then the transducer moved 90 degrees so that the needle was in-plane with the transducer using an ultrasound gel wedge to assist with in-plane positioning (Figure 3). The needle was then introduced through the joint capsule into the glenohumeral synovial space and lidocaine injected to confirm needle positioning (Figure 4). After injection of the lidocaine, arthrocentesis was performed and joint effusion was aspirated if present. The syringe was then detached from the needle while still in the glenohumeral joint and a 3 ml syringe prefilled with 60 mg triamcinolone acetonide suspension was attached to the indwelling needle, and the treatment was slowly injected. If there were unexpected resistance to injection, needle tip positioning was examined with ultrasound and the needle was rotated to change bevel positioning until resistance subsided and flow into the joint space could be confirmed, and the remainder of treatment was injected. The needle was then extracted, firm pressure applied to the puncture site, and a sterile bandage was applied.

**Figure 2.**
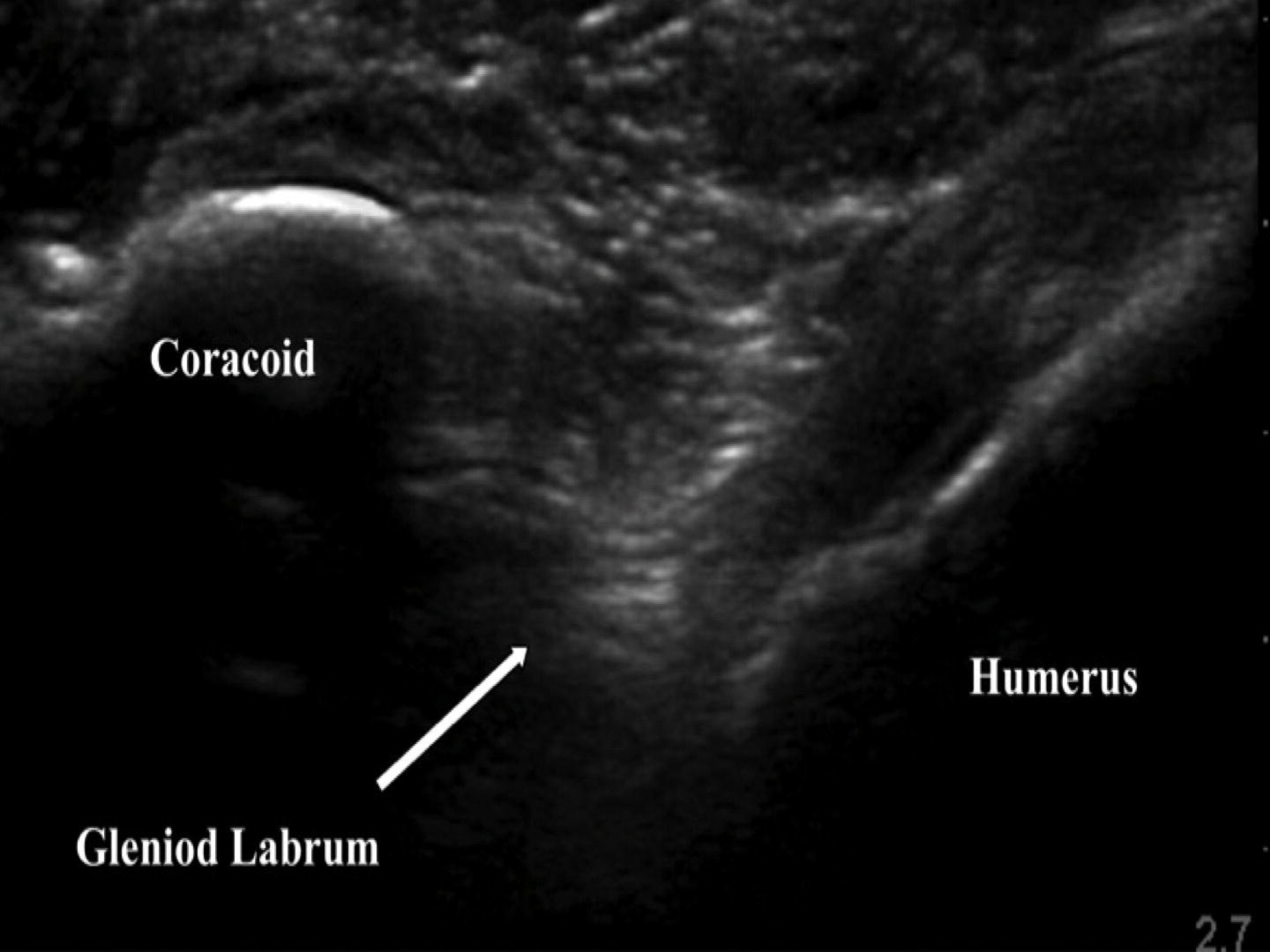
Ultrasound identification of the glenohumeral joint. In the ultrasound-guided technique the linear probe is held horizontally and the coracoid process and humerus are identified. The glenoid labrum and the glenohumeral joint are deep and between these structures. The needle is introduced out of plane to the probe inferiorly and laterally to the coracoid and medially to the humerus.

**Figure 3.**
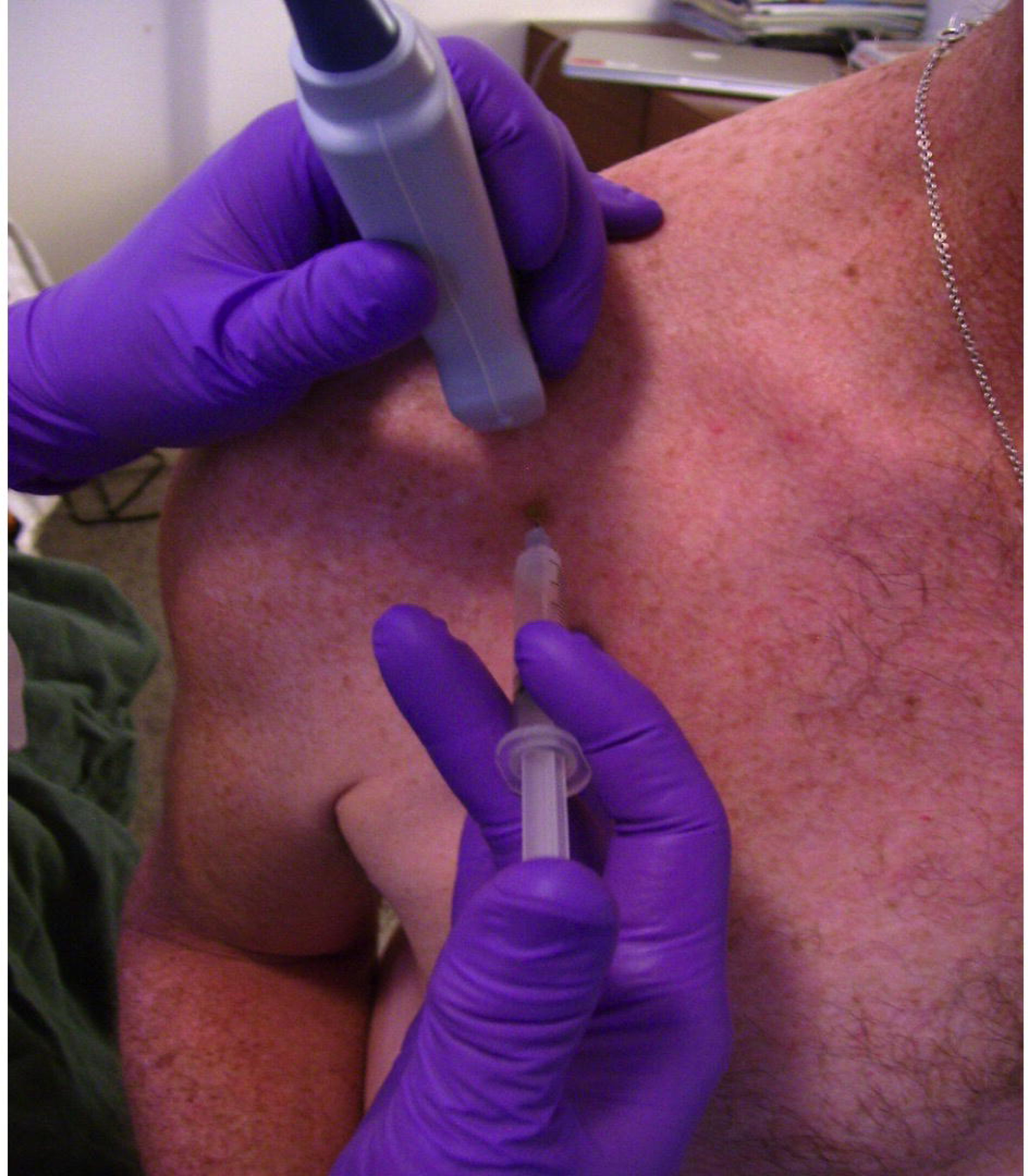
Probe and needle positioning. The needle is introduced out-of-plane inferiorly and laterally to the coracoid and then the transducer moved 90 degrees so that the needle is in-plane with the transducer using an ultrasound gel wedge to assist with in-plane positioning.

**Figure 4.**
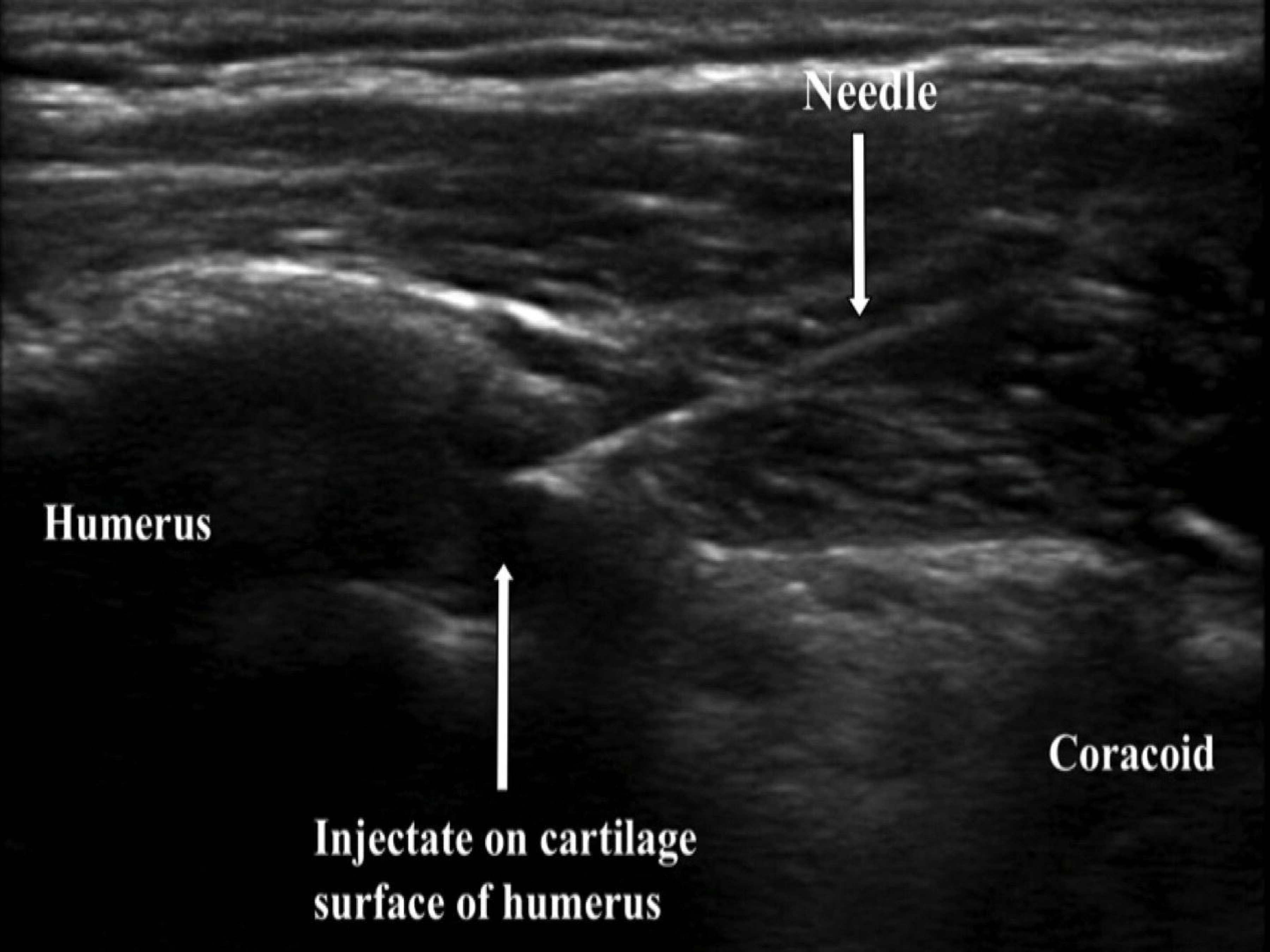
Injection of the glenohumeral joint. The needle is introduced through the joint capsule into the glenohumeral synovial space and lidocaine injected to confirm needle positioning. After injection of the lidocaine, the empty lidocaine syringe is detached from the needle while still in the glenohumeral joint and a 3 ml syringe prefilled with 60 mg triamcinolone acetonide suspension was attached to the indwelling needle, and the triamcinolone is slowly injected.

### Statistical Analysis

In joints other than the shoulder, ultrasound guidance has shown benefit in increasing the time-to-next-intervention and therapeutic duration, and thus, therapeutic duration was the primary outcome measure in this study and was used for power calculations (18,19). A power calculation for the primary outcome of therapeutic duration (time after the injection when VAS rises to ≥ 5 cm) was made using preliminary data where α=5%, power = 0.8, and allocation ratio = 1.0 indicated that n=10 in each group would provide statistical power at the p<0.05 level. Data were entered into Excel (Version 5, Microsoft, Seattle, WA), and analyzed in SAS (SAS/STAT Software, Release 6.11, Cary, NC). Since the hypothesis was that ultrasound-guidance would be superior to anatomic landmark palpation-guided procedures and prolong therapeutic duration, the differences between parametric two group data were determined with the one tailed t-test with significance reported at the p <0.05 level.

## Results

The study groups were similar in age, gender, and baseline pain (Table 1). There were no complications, including no infections, patient injuries, nerve injuries, vascular complications, or unintended needle-sticks in this trial. Direct comparisons between the palpation-guided injections and sonographic guidance are shown in Tables 1 and 2. As can be seen in Table 1, there was no significant difference in pre-procedure pain between the two treatment groups (p =0.86). However, shoulder injection with sonographic guidance resulted in 26% less needle introduction pain (p=0.18) and 79% less injection pain (p=0.05) by VAS than conventional anatomic landmark guided shoulder injection. In patients receiving sonographic-guided injections there were no patients reporting significant pain (VAS > 5 cm) with either needle introduction or medication injection. In contrast, 3 patients reported significant needle introduction pain and 1 patient reported significant medication injection pain with conventional palpation guided injection. Thus, shoulder injection with sonographic guidance was significantly less painful than conventional injection.

**Table 2.** Outcomes and Costs of Glenohumeral Injection

|  | **Landmark Guidance** | **Sonographic Guidance** |  |  |  |
| --- | --- | --- | --- | --- | --- |
| **Number of Shoulders** | **15** | **15** | **Percent Difference** | **95% CI of difference** | **P Value** |
| **Pre-Procedure Baseline Pain (VAS Pain Scale)\*** | 7.9±1.3 | 8.0±1.8 | 1.3% | -1.2 <-0.1< 1.0 | 0.86 |
| **Pain at Outcome (2 weeks) (10 cm VAS)\*** | 1.8±2.7 cm | 2.2±2.4 cm | +6.8% | -2.2 <-0.4< 1.4 | 0.66 |
| **Reduction in Pain from Baseline (10 cm VAS)\*\*** | 6.1±2.4 cm (-77% from baseline, p<0.001) | 5.8±2.5 cm (-73% from baseline, p<0.001) | -5% | -1.5 <0.3< 2.1 | 0.37 |
| **Responders (10 cm VAS < 2 cm)\*\*** | 53% (8/15) | 53% (8/15) | 0% | Not applicable | 0.28 |
| **Non-Responders (10 cm VAS ≥ 2 cm)\*\*** | 47% (7/15) | 47% (7/15) | 0% | Not applicable | 0.28 |
| **Pain at Outcome (6 months) (10 cm VAS)\*** | 6.4±2.9 cm | 5.8±2.8 cm | -9.1% | -0.4 <0.6< 1.1 | 0.71 |
| **Duration of Therapeutic Effect VAS≥5 (months)\*** | 3.0±1.2 months | 3.9±1.5 months | +30% | -1.4 <-0.9< -0.4 | 0.045 |
| **Time to Next Intervention** | 5.7±2.9 months | 8.1±3.5 months | +42% | -3.6 <-2.4< -1.3 | 0.025 |
| **Cost/Patient/Year – Physician Office (\$US)\*** | \$301±67 | \$318±89 | +5.6% (+\$17) | -45 <-17< 11 | 0.28 |
| **Cost/Responder/Year – Physician Office (\$US)\*** | \$564±141 | \$596±97 | -5.6% (+\$32) | -75 <-32< 11 | 0.23 |

\*Mean± Standard Deviation, one-tail t-test
\*\*Fisher Exact test

Intermediate-term (2 week) and long-term (6 month) therapeutic responses to the two injection methods are shown in Table 2. Sonographic-guided and landmark-guided injections were very similar in terms of pain at 2 weeks (Sonographic: 2.2±2.4 cm; Landmark: 1.8±2.7 cm; 95% CI of difference: −2.2 <−0.4< 1.4, p=0.66) and pain at 6 months (Sonographic: 5.8±2.8 cm; Landmark: 6.4±2.9 cm; 95% CI of difference: −0.4 <0.6< 1.1, p=0.71), indicating almost identical outcomes (Table 2).

However, sonographic guidance resulted in a 30% prolongation of therapeutic effect (the time interval when the VAS≥**5**) (Sonographic: 3.9±1.5 months; Landmark: 3.0±1.2 months; 95% CI of difference: −1.4 <−0.9< −0.4, p=0.045) and a 42% increase in the time-to-next-intervention (Sonographic: 8.1±3.5 months; Landmark: 5.7±2.9 months; 95% CI of difference: −3.6 <−2.4< − 1.3, p=0.025) (Table 2). Ultrasound guidance resulted in a 29% reduction in the number of corticosteroid injections required per year: Sonographic: 1.5±0.2 injections/year compared to Landmark: 2.1±0.2 injections/year (p<0.037; 95% CI of difference −0.9 <−0.6< −0.3). Thus, although both techniques were equally effective in terms use of relief of shoulder pain at 2 weeks, sonographic guidance was associated with a longer therapeutic duration, an extended time-to-next-intervention, and fewer corticosteroid injections per year.

Procedural costs for a 3^rd^ party payer are shown in Table 2 (US Medicare). The use of sonographic guidance in a physician office for injecting the shoulder modestly increased the costs/patient/year by 5.6% ($17) (p=0.28) and cost/responder/year by 5.6% ($32) (p=0.23) relative to palpation-guided landmark methods.

## Discussion

The present study confirms prior reports that shoulder injection therapy with anatomic landmark palpation guidance and ultrasound guidance are both effective for relieving chronic shoulder pain due to osteoarthritis (1-3,5,6). As has been reported previously in other joints, ultrasound-guidance increased the therapeutic duration and the time-to-next-intervention, and reduced the number of corticosteroid injections per year (18,19). However, despite these benefits, ultrasound-guidance was not statistically more effective at reducing pain at 2 weeks and 6 months, and in a patient with a painful osteoarthritic shoulder, modestly increased the costs/patient/year and costs/responder/year (Table 2).

A number of studies have supported the belief that sonographic guidance provides greater accuracy and improved outcomes for injection procedures (19,26,27). Pendelton et al, Raza et al, and Im et al demonstrated that sonography improves injection accuracy, and Qvistgaard et al showed that sonography can predict the response to corticosteroid injection (26-29). Eustace et al reported that patients whose injections had been accurately placed into the shoulder improved to a greater degree in the short term than those whose injections had been less accurately placed (30). In a similarly powered study to the present study, Naredo et al reported that US improved the 6-week outcomes of subacromial shoulder injections (31).

Thus, the literature indicates that sonographic guidance improves accuracy and outcomes of musculoskeletal injections, but outcomes and cost-effectiveness in specific procedures has not been addressed to the satisfaction of many (10-12). In the present study, image-guided shoulder injections were significantly less painful than landmark-guidance, causing 79% less absolute injection pain and 100% less significant pain (Table 1). The beneficial reduction in procedural and injection pain with ultrasound guidance has been reported previously and may be due to better control and direction of the needle tip away from pain-sensitive structures (18,19,32). An alternative explanation is that the cooling effect of ultrasound gel, the pressure from the ultrasound transducer, and a distracting effect at the neurocognitive level all significantly reduce anxiety and pain (18,19).

Mattie et al have recently demonstrated that inexperienced physicians using anatomic landmark guidance have only a 37% accuracy of injecting into the glenohumeral joint while experienced proceduralist have an accuracy of 64.4% (8). Simoni et al performed a review of the literature with a total 2690 injections of the shoulders and found that accuracy of glenohumeral injections varied from 42% to 100% and that image guided injections were on average more accurate than anatomic landmark injections (5). A recent review by Aly et al of published studies of over 800 shoulders confirmed improved accuracy with ultrasound in shoulder injections, including the glenohumeral joint with greater reduction in pain scores by VAS, a finding the present study did not confirm (33).

The present study demonstrated that in terms of pain scores at 2 weeks and 6 months, there was no difference between the anatomic landmark group and the ultrasound-guided injection group (Tables 1 and 2). Raeissadat et al have recently demonstrated that US improves the accuracy of glenohumeral injections but clinical pain parameters did not statistically improve, similar to the present study (6). Also, Cole et al recently demonstrated that US-guided subacromial injections were not superior to anatomic landmark injections in the shoulder (34). Sidon et al reported that anterior injections of the glenohumeral joint by anatomic landmark guidance has 98% accuracy which is comparable to ultrasound or fluoroscopy, thus, differences in injection outcome would not be expected (21). Similarly, Jo et al reported 91% accuracy and Kraeulter et al reported 93% accuracy injecting the glenohumeral joint using anatomic landmark markings (22,23). Further, Tobola et al found the anterior approach to glenohumeral injections as used in the present study to be more accurate than the posterior or supraclavicular approaches (24).

Hegedus et al found that accuracy in general had very little to do with outcome in glenohumeral injections and that extra-articular near-hit injections were as effective as accurate glenohumeral injections (35). Similarly, Bloom and Buchbinder and colleagues have recently reviewed the literature regarding ultrasound-guided shoulder injections and found that although ultrasound-guided injections are more accurate, that injection into a distant region is equally effective suggesting that a non-intraarticular systemic or regional lymphatic spread of the steroid is equally effective (36). Further, Dogu et al found that ultrasound -guidance and anatomic landmark injections of the subacromial joint result in identical outcomes, similar to the present report in the glenohumeral joint (37). Ekeberg et al demonstrated no difference in reduction of pain and disability in subacromial shoulder pain after 6 weeks in groups injected in the subacromial space with US versus a group getting gluteal injection (38). Oh et al reported the same result for frozen shoulder with US guided injection in the subacromial space versus the glenohumeral joint (39). It is known that once corticosteroid crystals are injected in the soft tissues in the proximity to an intended injection target that the corticosteroid can move via lymphatics or systemically, which may explain why exact accuracy is not always necessary (40).

Sage et al have recently reviewed the literature on the effects of US guidance vs. anatomic landmark injection and have found some consistent improvements, but these improvements are very small and that cost-effectiveness studies are required (41). In this line, the present study preliminarily determined the costs of glenohumeral injection with sonographic guidance (Table 2). Cost-effectiveness in these narrow domains is dependent on balancing the increased costs of sonographic guidance with a reduced use of other healthcare resources, in this case, reducing the costs of the next injection, primarily by prolonging the time-to-next-intervention (15-19). Although sonographically directed procedures were superior in therapeutic duration and the time-to-next-intervention (Table 2), sonographic image guidance for injection of the shoulder increased the costs per patient per year by 5.6% ($17), thus, ***modestly decreasing*** cost-effectiveness (Table 2) (10-12, 20). However, this difference is cost is rather small, and may not be significant if other health benefits are associated with ultrasound guidance.

Although intraarticular corticosteroids have beneficial short-term effects on arthritis pain, frequent exposure have been shown to damage the articular cartilage (42,43). McAlindon et al used an intensive regimen of triamcinolone injections administered intraarticularly into the knee every 3 months for 2 years and found that this regimen resulted in significantly more cartilage loss than control saline injections, a finding that has been confirmed in subsequent epidemiologic studies (43,44). These same studies have not been completed in the shoulder to date, but are likely a to be similar (42). Thus, a technique like ultrasound guidance that might result in fewer corticosteroid injections per year might mitigate these deleterious corticosteroid effects on articular cartilage (42-46).

There are recognizable limitations to the above analyses. Firstly, the study was relatively small with 30 shoulders, and all patients in the study were older female adults owing to the demographics of this clinical population. Power calculations indicated that the same results would be obtained at 2 weeks if the total population were increased to over 1000 shoulders, again emphasizing how similar the responses were between ultrasound-guided and landmark-guided techniques. The above analyses concerning the glenohumeral joint do not apply to other joints, since the anatomy and injection target characteristics are quite different (26-31). Also, the glenohumeral joint can be injected with an anterior, posterior, or superior approach with variable accuracy, thus, the present research only relates to anterior glenohumeral injections (47-49). Subacromial injections also are quite distinct so the present research does not apply to those injections (7, 31,34,37-39). Physician or institutional costs including the expense of acquisition and maintenance of the ultrasound machine, image storage, and sonographic supplies, the increased operator set up and procedure time were not included in this analysis, but if included would tend to further degrade cost-effectiveness of ultrasound guidance.

## Conclusions

Anatomic landmark guidance for corticosteroid injection of the painful osteoarthritic shoulder in the short-term is equally effective as ultrasound and modestly less costly, however, ultrasound guidance may reduce the need for repetitive injections by prolonging the time-to-next-injection.

## Acknowledgment of grants, disclosures, or other assistance

None of the authors (Moore, Paffett, Sibbitt, Hayward, Fields, Emil, Fangthm) have any grants, disclosures, or other assistance to report.

There was no industry support for this study.

## Disclosures

None of the authors declare conflict of interest.

